# Mycophenolate mofetil increases inflammation resolution in zebrafish via neutrophil apoptosis

**DOI:** 10.1101/145359

**Authors:** Aleksandra Bojarczuk, Simon A. Johnston

## Abstract

Mycophenolate mofetil (MMF) is an immunosuppressive agent used in the treatment of autoimmune and inflammatory conditions, and following organ transplant. MMF treatment results in lymphopenia via the depletion of purines required for DNA synthesis. While the primary effect of MMF treatment is thought to be via the depletion of lymphocytes, MMF has also been associated with innate immune defects, including neutropenia and neutrophil dysplasia. Here, we address the question of MMF specific effects on neutrophils in an *in vivo* model of neutrophil inflammation in zebrafish. We find that, following tissue injury, MMF increases resolution of neutrophilic inflammation via increased neutrophil apoptosis. Critically, we identify that the effect of MMF is distinct from DNA synthesis inhibition by using the competitive inhibitor of purine nucleotide incorporation, azathioprine. Therefore, we propose that increased neutrophil cell death during inflammatory insult may play a role in neutrophil defects associated with MMF treatment.

## Introduction

Mycophenolate mofetil (MMF) is a commonly prescribed immunosuppressive used as an anti-proliferative agent against lymphoid cells. MMF acts by inhibiting the inosine monophosphate dehydrogenase (IMPDH) enzyme, which is critical in the production of purine nucleotides. Due to the differential requirements for IMPDH among immune cell types, MMF treatment induces lymphopenia by depleting purine nucleosides available for DNA synthesis preferentially in lymphoid cells. While the mechanism of MMF action is distinct to lymphoid cells, MMF treatment is recognised to cause neutropenia in some patients (Shapiro et al., 1999). In addition, there are several cases of MMF causing neutrophil dysplasia (Banerjee et al., 2000).

Neutrophils are essential in host defence against pathogens and are rapidly recruited to sites of infection and injury. The resolution of inflammation requires the clearance of neutrophils (Savill et al., 1989). Failure in this clearance causes chronic inflammation and is an underlying pathology of many diseases. However, in the absence of neutrophils the host is profoundly immunocompromised and highly susceptible to infection (Estcourt et al., 2015). Resolution of neutrophil resolution can occur via neutrophil apoptosis, and clearance by macrophages, and/or by reverse migration of neutrophils out of inflamed tissue (Nourshargh et al., 2016; Robb et al., 2016).

Azathioprine (AZA) is a second lymphoid cell anti-proliferative agent that is both an alternative to MMF and associated with neutropenia. Clinical studies of MMF versus AZA have associated MMF with lower incidence of leukopenic adverse effects (Eisen et al., 2005). However, while neutropenia with AZA treatment is associated with several genetic predispositions (Coulthard et al., 2004), the mechanism of MMF associated neutropenia is unknown. Several lines of evidence suggested to us that inflammatory insult might be a defining factor in neutropenia in MMF patients: There is underlying inflammation in several patient groups that receive MMF (e.g. liver transplantation following chronic hepatitis) (Nogueras et al., 2005), neutrophils from a patient following a kidney transplant showed an enhanced oxidative burst with MMF treatment (Hochegger et al., 2006), and that MMF alone has been shown to be insufficient to cause a neutrophil defect (Etzell and Wang, 2006).

Therefore, we hypothesised that MMF treatment caused a defect in neutrophil function in inflammation resolution, either due to increased neutrophil apoptosis or a defect in reverse migration. To address this hypothesis, we used the well-described zebrafish model of neutrophilic inflammation. We identified that there were reduced numbers of neutrophils with MMF treatment following tissue injury. AZA had no effect on total neutrophil numbers or inflammation resolution. We show that increased resolution of neutrophilic inflammation is driven by neutrophil apoptosis with no change in reverse migration.

## Results

### Neutrophilic inflammation is reduced with MMF treatment

Using a zebrafish tail transection assay of neutrophilic inflammation (Fig. 1A), where neutrophils are recruited to the injury site and, from ~6 hours post wounding, inflammation resolves, we found that MMF reduced the number of neutrophils at the wound site in the resolution stage of inflammation (Fig1. B). Increased resolution was indistinguishable from treatment with the positive control, the c-Jun N-terminal kinase inhibitor, SP600125 (Fig. 1B) (Robertson et al., 2014). Counting the number of neutrophils in the recruitment phase of inflammation demonstrated that there was also a small but significant reduction in the number of neutrophils recruited to the site of injury in MMF treated embryos (Fig. 1C). Our positive control, inhibition of c-Jun N-terminal kinase, had a larger effect on recruitment compared to MMF (Fig. 1C; Median DMSO=26, SP600125=14.5, MMF=20). To find the relative contribution of recruitment and resolution in the decrease neutrophilic inflammation with MMF treatment we calculated the percentage change in the number of neutrophils at the site of injury in the same zebrafish between 6 and 24 hours. We found that MMF treatment resulted in a more than 2-fold reduction in the proportion of neutrophils at the site of injury between 6 and 24 hours (Fig. 1D; Median DMSO=28%, MMF=64%), evidence that reduced inflammation with MMF treatment was due to increased resolution. Reduced recruitment of neutrophils may have been due to a smaller total number of neutrophils with MMF treatment. To test this possibility, we took advantage of the ability to image throughout the zebrafish larvae, counted the total number of neutrophils in zebrafish larvae following treatment for 24 hours with MMF. We found that there was no difference in the total number of neutrophils compared to control treatment (Fig. 1E–G). To understand the specificity of the effect of MMF treatment, we examined the related anti-proliferative immunosuppressant azathioprine (AZA).

**Figure 1.**
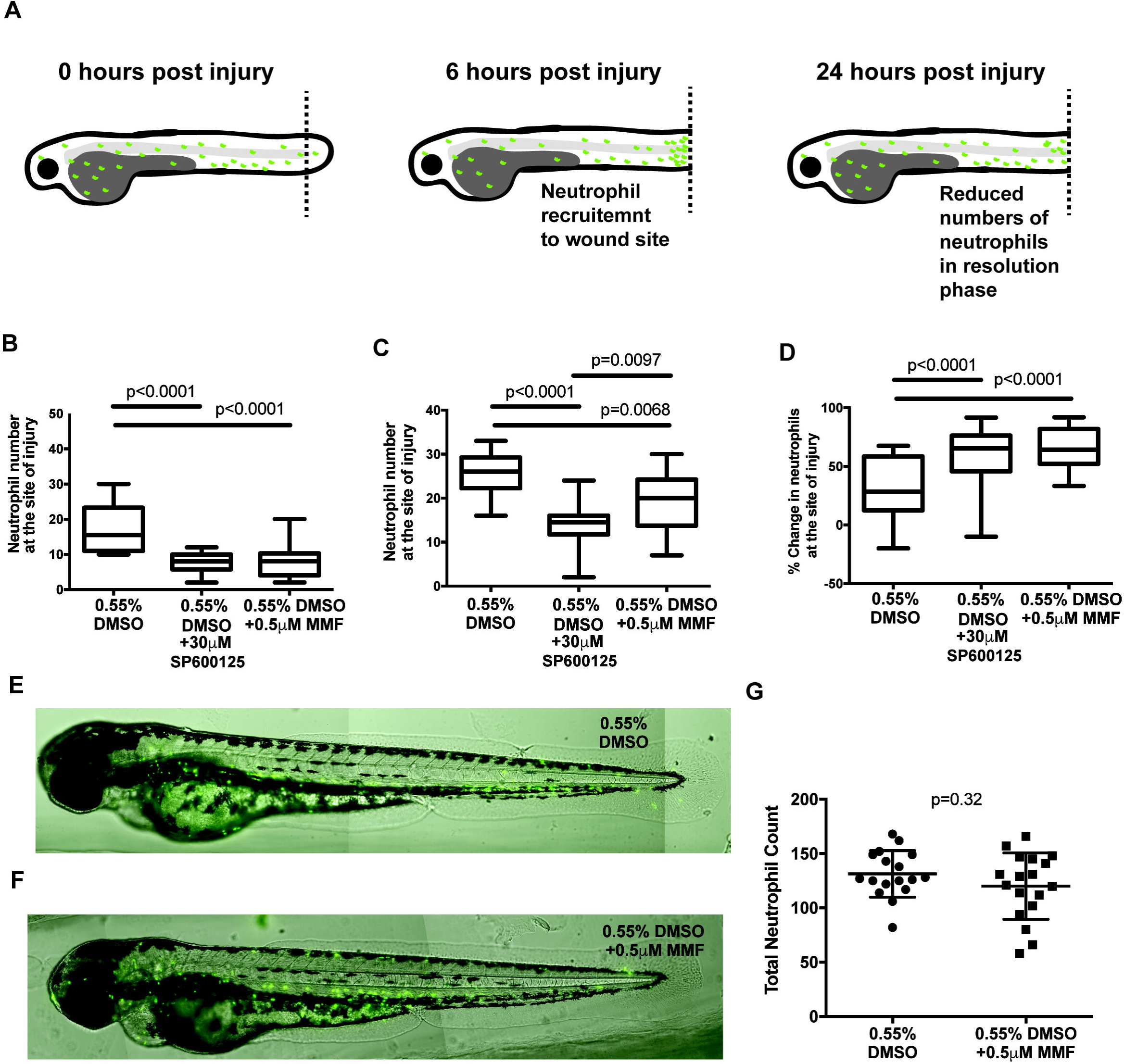
Mycophenolate mofetil reduces the number of neutrophils at the site of injury. A. Zebrafish model of neutrophilic inflammation. Tails of 3 day post fertilisation larvae were cut and the number of neutrophils at the site of injury counted at 6 hours (recruitment phase) and 24 hours (resolution phase). B-G *Tg(mpx:GFP)i114* 18 animals per group, 6 per 3 biological repeats. B. Number of neutrophils at the site of injury at 24 hours post injury (hpi). C. Number of neutrophils at the site of injury at 6hpi. D. Percentage change in number of neutrophils at the site of injury calculated by dividing the number of neutrophils at 6hpi by the number at 24hpi expressed as a percentage. E,F. Representative images of zebrafish treated with MMF or vehicle for 24 hours. G. Total number of neutrophils with MMF or vehicle treatment. All p-values are Mann-Whitney U test.

### Azathioprine does not recapitulate the effect of mycophenolate mofetil treatment on neutrophilic inflammation

In contrast to MMF, AZA had no effect on either recruitment or resolution of neutrophils at the site of injury (Fig. 2A,B). The mechanism of AZA is distinct from MMF as, instead of inhibiting the production of guanine nucleosides, AZA generates 6-thioguanidine that blocks nucleoside addition during nucleic acid synthesis. In addition, we found that 24 hour treatment with AZA had no effect on the total number of neutrophils (Fig 2.C).

**Figure 2.**
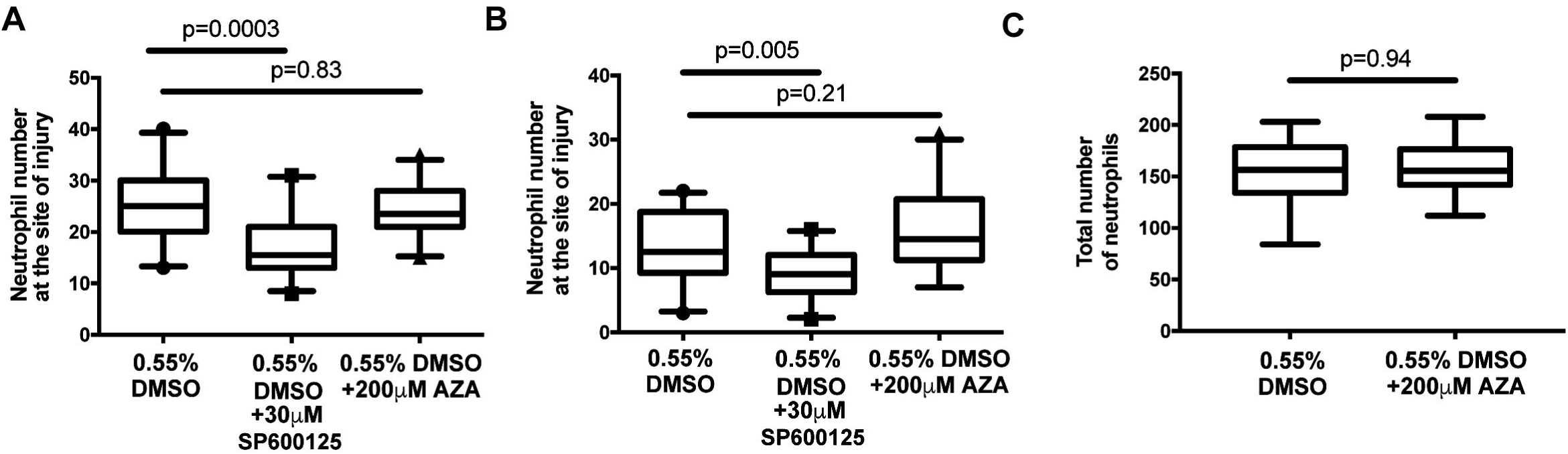
Azathioprine does not affect neutrophil recruitment, resolution or total numbers in a zebrafish model of neutrophilic inflammation. A-C *Tg(mpx:GFP)i114* 18 animals per group, 6 per 3 biological repeats. A. Number of neutrophils at the site of injury at 24 hours post injury (hpi). B. Number of neutrophils at the site of injury at 6hpi. C. Total number of neutrophils with AZA or vehicle treatment. All p-values are Mann-Whitney U test.

### MMF does not increase reverse migration of neutrophils from the site of injury

Using florescence time lapse imaging of photo converted neutrophils at the site of injury (Fig. 3A) we found that MMF did not alter reverse migration of neutrophils from the site of injury (Fig. 3B–F). Therefore, while we had shown there were different numbers of neutrophils at the site of injury with MMF, there was no difference in the number or rate of neutrophils leaving the site of injury (Fig. 3B). Given there was no difference in reverse migration we tested if there was increased apoptosis at the site of injury as the mechanism of increased neutrophil inflammation.

**Figure 3.**
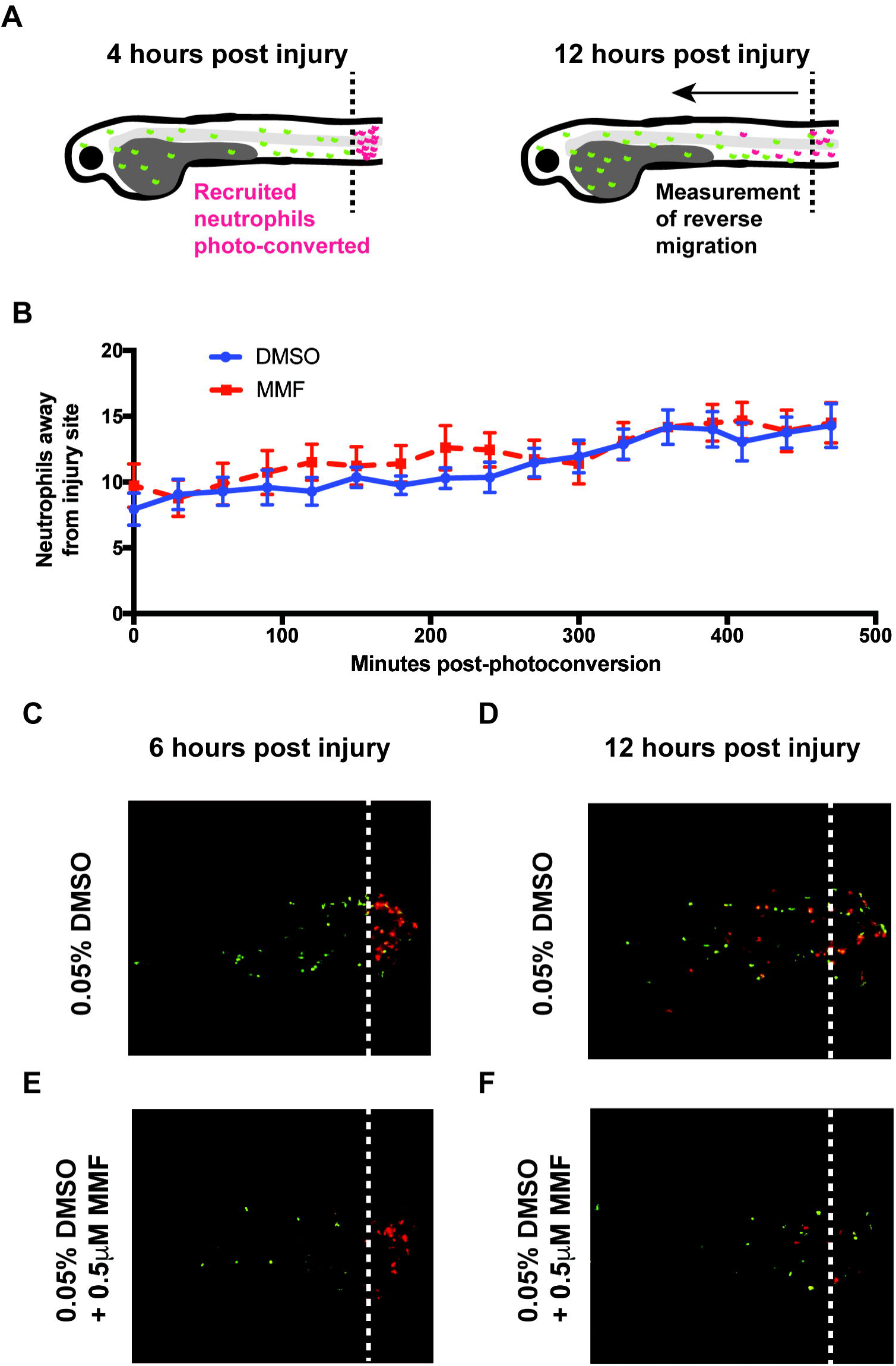
MMF does not alter reverse migration of neutrophils from the site of injury. A. A zebrafish model of neutrophil reverse migration. 4 hours post tail injury neutrophils at the site of injury are photoconverted (from green to red) for 2 hours and imaged from 6 to 12 hours. Reverse migrated neutrophils are red neutrophils that leave the area of photoconversion. B. The number of reverse migrated neutrophils was counted every thirty minutes over 6 hours for MMF and vehicle treatment. Points are the mean and the standard deviation of 17 (vehicle) or 18 (MMF) animals from three biological repeats. Difference in slope was assessed by linear regression analysis with p=0.52 chance of the lines being different by chance. C-F. Images of zebrafish at 6 and 12 hours post injury, following photoconversion.

### MMF increases neutrophil apoptosis at the site of inflammation

We used TUNEL staining *in vivo* to identify apoptotic cells at 12 hours post injury (hpi), an intermediate stage of inflammation resolution (Fig.4 A,B). At 12 hours hpi there was a significant difference in the number of neutrophils at the site of injury of MMF treatment (Fig. 4C). We counted the number apoptotic neutrophils (double *mpx*:GFP and TUNEL positive cells) and identified a >2-fold increase in neutrophil apoptosis with MMF treatment (Fig. 4D). Finally, we tested if inhibition of neutrophil cell death was sufficient to reverse the increased inflammation resolution we had identified with MMF. Using the pan-caspase inhibitor Z-VAD to inhibit apoptosis, we found that we could reverse the effect of MMF on the number of neutrophils at both 6 and 24 hours, demonstrating that preventing neutrophil cell death was sufficient to reverse neutrophilic inflammation resolution due to MMF (Fig. 4E,F).

**Figure 4.**
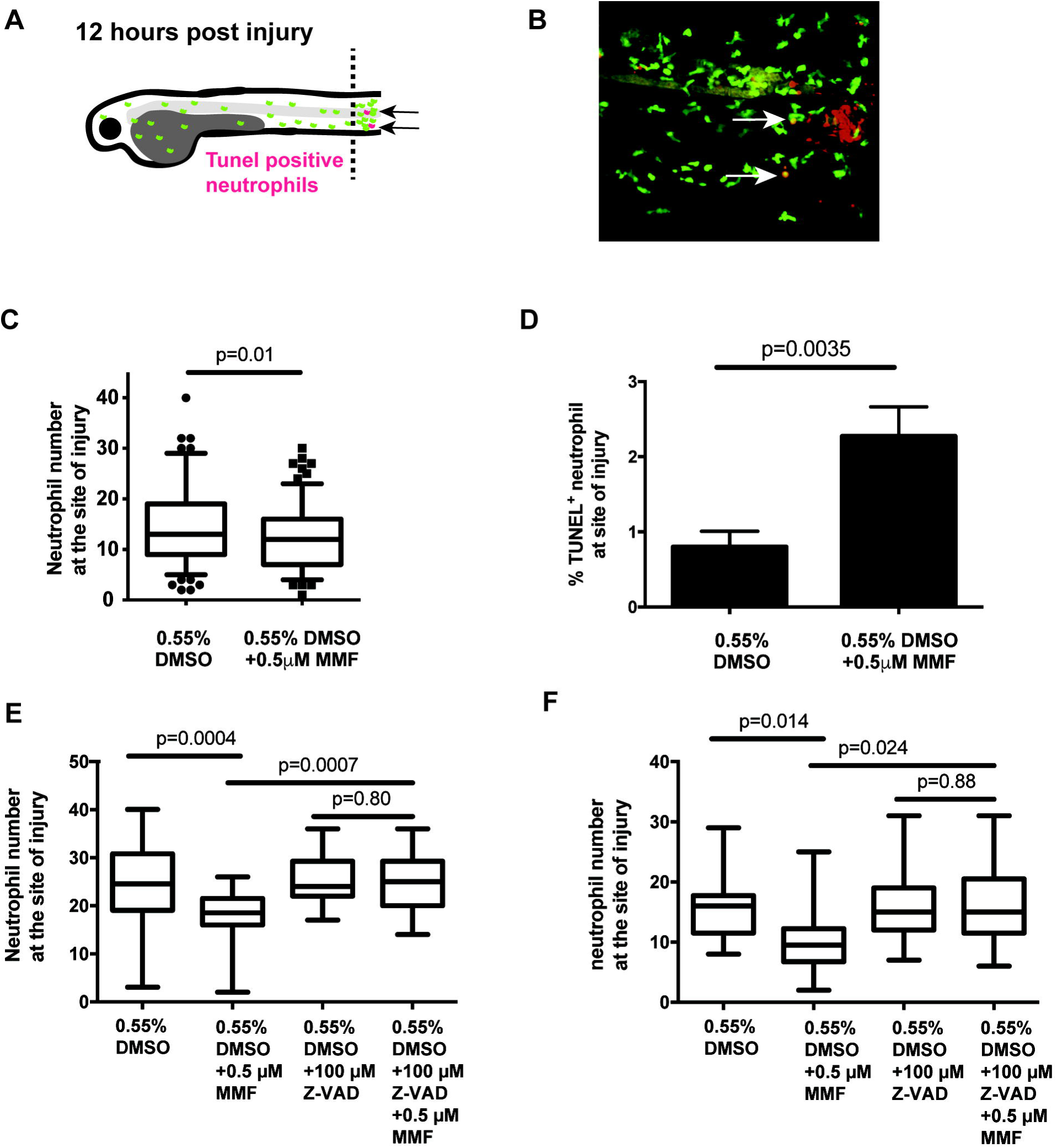
MMF increases resolution of neutrophilic inflammation via increased apoptosis at the site of injury. A. Measurement of neutrophil apoptosis with TUNEL staining at 12 hours post injury, an intermediate time of inflammation resolution. B. Image of *Tg(mpx:GFP)i114* zebrafish stained for TUNEL at 12 hours post injury, neutrophils, green and TUNEL stain, red. Arrows, apoptotic neutrophils yellow, dual stained with GFP and TUNEL. C. Quantitation of the number of neutrophils at the site of injury 12 hours post injury. B, C. 140 vehicle treated and 161 MMF treated animals from three biological repeats D. Quantitation of the percentage of apoptotic neutrophils at the site injury 12 hours post injury. E. Number of neutrophils at the site of injury at 24 hours post injury (hpi) with inhibition of apoptosis with the caspase inhibitor Z-VAD. F. Number of neutrophils at the site of injury at 6 hours post injury (hpi) with inhibition of apoptosis with the caspase inhibitor Z-VAD. E, F. *Tg(mpx:GFP)i114* 18 animals per group, 6 per 3 biological repeats.

### Discussion

Here we have shown how the common immunosuppressive mycophenolate mofetil (MMF) treatment results in neutrophil cell death by apoptosis *in vivo*, thereby reducing neutrophilic inflammation. We initiated this study to investigate possible mechanisms of MMF induced neutropenia and our data suggests that there may be an inflammatory component in the neutropenia observed in MMF treated patients.

While a large proportion of patients receiving MMF have inflammatory disease, be it prior to organ transplant, as with chronic hepatitis, or as their primary treatment criteria, e.g. interstitial lung disease (Nogueras et al., 2005; Volkmann et al., 2017), to our knowledge, this is the first time that an association between MMF associated neutropenia and inflammatory insult has been investigated. A clinical study of a patient with acute inflammatory syndrome, following kidney transplantation and MMF treatment, identified an increased neutrophil oxidative burst but only following phorbol myristate acetate and formyl Met-Leu-Phe stimulation (Hochegger et al., 2006) supporting our conclusion that MMF treatment can enhance neutrophil inflammatory responses, including apoptosis.

We tested if thiopurines (generated by AZA) were also able to increase neutrophil inflammation following inflammatory insult. We found no effect on total neutrophil numbers, the recruitment or resolution of neutrophils at the site of injury. More than 20% of patients have AZA discontinued because of adverse events, most commonly due to bone marrow suppression induced leukopenia (Kim et al., 2016). Our data suggest that AZA and MMF induced neutrophil defects have distinct mechanisms. The mechanism of AZA induced leukopenia is not known in all cases but there is a recognised host genetic component with several polymorphisms identified (Colombel et al., 2000; Kim et al., 2016; Weinshilboum and Sladek, 1980; Yang et al., 2014). AZA is a prodrug that is metabolised both enzymatically and non-enzymatically to form thioguanine nucleotides. The polymorphisms associated with AZA neutropenia are in genes that regulate thiopurine deactivation, metabolism and repair of thiopurine mediated DNA damage (Colombel et al., 2000; Kim et al., 2016; Weinshilboum and Sladek, 1980; Yang et al., 2014).

MMF treatment decreased the numbers of neutrophils at the site of injury in the resolution phase but we identified that MMF treatment also resulted in reduced neutrophil numbers at 6 hours where recruitment was expected to peak. We have two possible explanations for this difference; given our finding that MMF increased neutrophil apoptosis it is possible that with MMF treatment apoptosis occurs earlier than normal and therefore there is already a reduction in neutrophils at the site of injury at 6 hours due to early induction of neutrophil cell death. Secondly, MMF has been reported to reduce levels of cytokine-induced neutrophil chemoattractant (CINC; CXCL1) in a rodent model of lung perfusion injury and therefore MMF may directly reduce neutrophil numbers at the site of injury (Farivar et al., 2005). There implications for circulating neutrophil numbers are as a marker of bone marrow recovery following insult, most often following transplant. MMF has been associated with slower recovery of neutrophil numbers in an experimental model of graft versus host disease. However, such studies do not account for MMF treatment having a direct effect on circulating neutrophil numbers by increasing tissue accumulation of neutrophils (due to defects in chemotaxis following extravasation) and increased apoptosis shown in our study. In either case, there may be increased stress on the recovering bone marrow. In our study, we have only studied the short-term effects of MMF on neutrophil numbers but it is possible that with longer treatment, as in immunosuppressive therapy, there may be an impact on total neutrophil numbers.

The occurrence of neutropenia with MMF is an adverse effect that results in a reduction of MMF dose or replacement with an alternative agent. However, MMF is used also for the treatment of inflammatory conditions with a neutrophilic component. MMF reduced the vascular permeability and inflammatory cell recruitment and persistence in experimental models of lung perfusion injury (Farivar et al., 2005). Recent analysis of the scleroderma lung studies demonstrated that MMF was associated with increased forced vital capacity (FVC) and reduced shortness of breath in Systemic Sclerosis-Related Interstitial Lung Disease (Volkmann et al., 2017). Finally, our data, and the clinical evidence for use of MMF in inflammatory disease, suggest that neutrophils may be a *de facto* target for MMF. Recently, methotrexate, another chemotherapy drug used as an anti-inflammatory agent, has been identified as new inhibitor of the JAK/STAT pathway (Thomas et al., 2015). Methotrexate is classically an inhibitor of the folate pathway, via dihydrofolate reductase inhibition, but its novel JAK/STATS activity may represent a significant proportion of its anti-inflammatory effect. Thus, our study demonstrates a specific effect of MMF on neutrophil apoptosis and highlights the need for better mechanistic understanding of MMF action specifically and immunosuppressive treatment in general.

## Materials and Methods

### Ethics statement

These studies have been subjected to guidelines and legislation set out in UK law in the Animals (Scientific Procedures) Act 1986, under Project License PPL 40/3574. Ethical approval was granted by the University of Sheffield Local Ethical Review Panel.

### Fish husbandry

We used the neutrophil marked transgenic *Tg(mpx:GFP)i114* and Tg(mpx:Gal4);Tg(UAS:Kaede)i222 (Robertson et al., 2014) for the reverse migration assay. Zebrafish strains were maintained per standard protocols and local animal welfare regulations. Adult fish were maintained on 14–10 hour light cycle at 28 °C in UK Home Office approved facilities in the Bateson Centre aquaria at the University of Sheffield.

### Compound treatment of zebrafish larvae

All stock solutions of compounds were dissolved in DMSO (Sigma-Aldrich, UK; all reagents from Sigma unless otherwise stated) with a final concentration of DMSO being 0.55%. Treatment was performed by adding compounds to E3 solution to achieve the desired concentration. We used 0.05 µM MMF (Generon Ltd, UK), 30 µM SP600125, 100 µM Z-VAD (Insight Biotechnology Ltd, UK) and 200 µM AZA.

### Microscopy

Confocal images were captured using a Ultraview Nipkow spinning disk confocal (Perkin Elmer) on an Olympus IX81 inverted microscope with 1000x1000 Hamamatsu 9100–50 front illuminated EMCCD camera, with 10x objective lens, using Volocity 6.3 (Perkin Elmer, UK). Wide field fluorescence images were captured using a Nikon Ti-E with with a CFI Plan Apochromat λ 10X, N.A.0.45 objective lens and using Intensilight fluorescent illumination with ET/sputtered series fluorescent filters 49002 (Chroma, Bellow Falls, VT, USA). Images were captured with a Neo sCMOS camera (Andor, Belfast, UK) and NIS Elements 391 software (Nikon, Richmond, UK).

## Inflammation assays

### Recruitment

To test recruitment larvae were injured at 2 dpf, by tail fin transection to initiate an inflammatory response involving recruitment of neutrophils to the site of injury (Renshaw et al., 2006). The number of fluorescent neutrophils was counted by eye at 6 hpi to detect changes in neutrophil numbers that could indicate recruitment inhibiting effects of MMF. Zebrafish were anesthetized by immersion in 4.2% tricaine in E3 prior the count. 6 fish were used for each treatment over 3 biological repeats.

### Resolution

To assess MMF activity in accelerating resolution of inflammation, larvae were injured at 2 dpf and at 4 hpi (Renshaw et al., 2006). The number of fluorescent neutrophils recruited to inflammation site was counted by eye at 6 hpi and 24 hpi (2 and 20 hours after addition of MMF, respectively). After 6hpi count larvae were recovered in fresh E3 and returned to a new plate. Zebrafish were anesthetized by immersion in 4.2% tricaine in E3 prior the counts. 6 fish were used for each group in 3 biological repeats.

### Total number of neutrophils

To evaluate total number of neutrophils, uninjured fish were incubated with compounds for 6 and 24 hours, and imaged at these time-points. Prior to each time-point, fish were anesthetized by immersion in E3 with 4.2% tricaine and aligned in 1% agar channels for imaging (Bojarczuk et al., 2016). Channels were made by adding 200μl of 1% agar in 4,2% tricaine in E3 into 96-well glass-bottomed plates (Porvair sciences, UK). Mounting moulds were cut in solidified agar using GelX4 tips (Geneflow, UK) and embryos were imaged on a Nikon Ti-E microscope fitted with a climate controlled incubation chamber (28 °C; Okolabs, Pozzuoli, Italy). After 6hpi imaging, larvae were recovered in fresh E3 and returned to a new numbered 96-well plate to be then imaged at 24 hpi.

### Apoptosis

The occurrence of apoptosis was observed by dual staining for neutrophil-specific MPX activity-TSA staining (TSA plus kit, Fluorescence Systems, Perkin Elmer Inc., Waltham, USA) and then for an apoptotic marker –TUNEL (Rhodamine Apoptag Red In Situ kit was from Millipore UK Ltd, UK). An inflammatory response was induced as in the resolution assay and at 12 hpi larvae were fixed with 4% PFA (Loynes et al., 2010). For fluorogenic peroxidase substrates Fluorescein-TSA staining fish were washed 3 times in PBS, incubated at 28°C for 10 min in the dark in a 1:50 dilution of FITC TSA in amplification diluent. Then larvae were washed 3 times in PBS, 10 mins on each and fixed again at room temperature 4% PFA for 20 min. For ApopTag Red *in situ* apoptosis detection staining larvae were washed for 3 times 5 min each in PBS, then left at room temperature in 10 µg/ml proteinase K for 90 min, washed and stained using the ApopTag Red *in situ* apoptosis detection kit. The samples were kept in 75% glycerol at –20°C. For imaging larvae were mounted with 75% glycerol on microscope slides (VWR International Ltd, UK) with cover glass (VWR International Ltd, UK) and imaged by confocal microscopy. Rates of apoptosis were assessed by the percentage of TSA-positive neutrophils labelled via TUNEL.

### Reverse migration

In reverse migration assay, an inflammatory response was initiated as in the resolution assay and at 6 hpi neutrophils were photoconverted at the site of injury site. Photoconversion of neutrophil-specific Kaede protein was performed by using confocal microscopy as previously described (Elks et al., 2011). Behaviour of inflammatory neutrophils was imaged for 8 hours. For imaging, zebrafish were mounted in 0.8% low melting agarose and in E3 fish water both containing tested drug or vehicle controls. Reverse migration was assessed by counting neutrophils every 30 minutes over the period 8 hours in the posterior and anterior region to the circulatory loop. The region of interest was at the anterior of the tail and this data set was extracted to plot the number of photoconverted neutrophils leaving the area of transection.

### Statistical analysis

Statistical analysis was performed as described in the results and figure legends. Graphing and statistical analysis was performed in Prism 7.0.

## Acknowledgements

We thank Stephen Renshaw and Catherine Loynes for their expertise in zebrafish inflammation and for discussion of data. We thank the Bateson Centre aquaria staff for their assistance with zebrafish husbandry. SAJ and AB were supported by Medical Research Council and Department for International Development Career Development Award Fellowship MR/J009156/1. SAJ was additionally supported by a Krebs Institute Fellowship, Medical Research Foundation grant R/140419 and Medical Research Council Centre grant (G0700091). Confocal microscopy was supported by an MRC grant (G0700091), a Wellcome Trust grant (GR077544AIA) and was carried out in the Wolfson Light Microscopy Facility. The authors declare no competing financial interests.

